# Purification of synchronized *E. coli* transcription elongation complexes by reversible immobilization on magnetic beads

**DOI:** 10.1101/2022.01.10.475675

**Authors:** Skyler L. Kelly, Courtney E. Szyjka, Eric J. Strobel

## Abstract

Synchronized transcription elongation complexes (TECs) are a fundamental tool for *in vitro* studies of transcription and RNA folding. Transcription elongation can be synchronized by omitting one or more NTPs from an *in vitro* transcription reaction so that RNA polymerase can only transcribe to the first occurrence of the omitted nucleotide(s) in the coding DNA strand. This approach was developed over four decades ago and has been applied extensively in biochemical investigations of RNA polymerase enzymes, but has not been optimized for RNA-centric assays. In this work, we describe the development of a system for isolating synchronized TECs from an *in vitro* transcription reaction. Our approach uses a custom 5’ leader sequence, called C3-SC1, to reversibly capture synchronized TECs on magnetic beads. We first show that complexes isolated by this procedure, called ^C3-SC1^TECs, are >95% pure, >98% active, highly synchronous (94% of complexes chase in <15s upon addition of saturating NTPs), and compatible with solid-phase transcription; the yield of this purification is ∼8%. We then show that ^C3-SC1^TECs perturb, but do not interfere with, the function of ZTP-sensing and ppGpp-sensing transcriptional riboswitches. For both riboswitches, transcription using ^C3-SC1^TECs improved the efficiency of transcription termination in the absence of ligand but did not inhibit ligand-induced transcription antitermination. Given these properties, ^C3-SC1^TECs will likely be useful for developing biochemical and biophysical RNA assays that require high-performance, quantitative bacterial *in vitro* transcription.

## Introduction

In the past several years, bacterial transcription systems have been used to develop high-resolution biochemical and biophysical methods for studying RNA folding and function (1-12). In many cases, biochemical tools that were developed for studying the process of bacterial transcription have been successfully repurposed for these applications. Among these tools, strategies for synchronizing transcription (13) are particularly important because they generate a virtually uniform population of transcription elongation complexes (TECs) that can be used in biochemical assays.

It is well-established that transcription complexes can be synchronized *in vitro* by ‘walking’ RNA polymerase (RNAP) to a synchronization site (13-15). In this approach, one or more NTPs are omitted from a transcription reaction so that RNAP transcribes until it reaches the first occurrence of the nucleotide that was omitted. Transcription then resumes when the NTP that was omitted is added to the transcription reaction. While transcription complexes are occasionally synchronized using natural DNA sequences, it is more common to initiate transcription using one of several established leader sequences that facilitate the formation of stable, synchronized TECs. This approach is applied routinely in high-resolution analyses of transcription elongation kinetics (13). For this application, the sequence of the 5’ leader and the purity of the synchronized TECs are typically inconsequential as long as the complexes are stable and resume transcription synchronously. However, when preparing synchronized TECs for assays that measure RNA folding or function, the sequence of the 5’ leader is crucial because it must be sequestered from interactions with the downstream RNA of interest. Furthermore, in some cases the ability to purify synchronized TECs may be advantageous because it establishes a quantitative relationship between template DNA molecules and their RNA product. A standardized system for generating high-purity synchronized TECs for use in RNA biochemical assays could therefore aid the development of new technology for investigating RNA structure and function.

In this work, we describe a procedure for isolating synchronized *E. coli* TECs from an *in vitro* transcription reaction using a 33 nt 5’ leader sequence, called C3-SC1. Our approach builds upon established procedures for synchronizing bacterial transcription by integrating an oligonucleotide hybridization site and a fast-folding RNA hairpin with an A-less cassette that can be used to walk RNAP to a synchronization site. Together, these elements facilitate the purification of synchronized TECs using a straightforward reversible immobilization procedure and minimize potential interactions between the C3-SC1 transcript and downstream RNA nucleotides by sequestering 31 of 33 5’ leader nucleotides. We first show that TECs isolated using the C3-SC1 leader (^C3- SC1^TECs) are >95% pure, >98% active, and highly synchronous (94% resume transcription in <15 s upon addition of saturating NTPs). We then show that two distinct transcriptional riboswitches remain biochemically active when fused to the C3-SC1 leader. Notably, appending the C3-SC1 leader to these riboswitches reduced basal transcription terminator readthrough but did not reduce the dynamic range of the ligand-induced transcription antitermination response. Overall, these findings establish a procedure for purifying synchronized TECs to homogeneity.

## Results

### Overview of the strategy for purifying synchronized TECs

The purpose of the procedure described below is to isolate synchronized *E. coli* TECs from an *in vitro* transcription reaction. Isolating synchronized TECs requires a strategy for separating TECs from both free DNA and promoter complexes that failed to initiate transcription. One fundamental difference between TECs and non-productive promoter complexes is that only the former contain nascent RNA. We therefore envisioned that the RNA product of an appropriately designed 5’ leader sequence could be used to purify synchronized TECs using a reversible immobilization strategy. In this approach, TECs are walked to a synchronization site and a capture oligonucleotide containing a photocleavable biotin modification (16) is annealed to nascent RNA (Figure 1A). TECs can then be isolated by immobilization on streptavidin-coated magnetic beads and eluted using 365 nm UV light (Figure 1A). The resulting complexes can then be used in downstream applications that require high-purity synchronized TECs.

**Figure 1.**
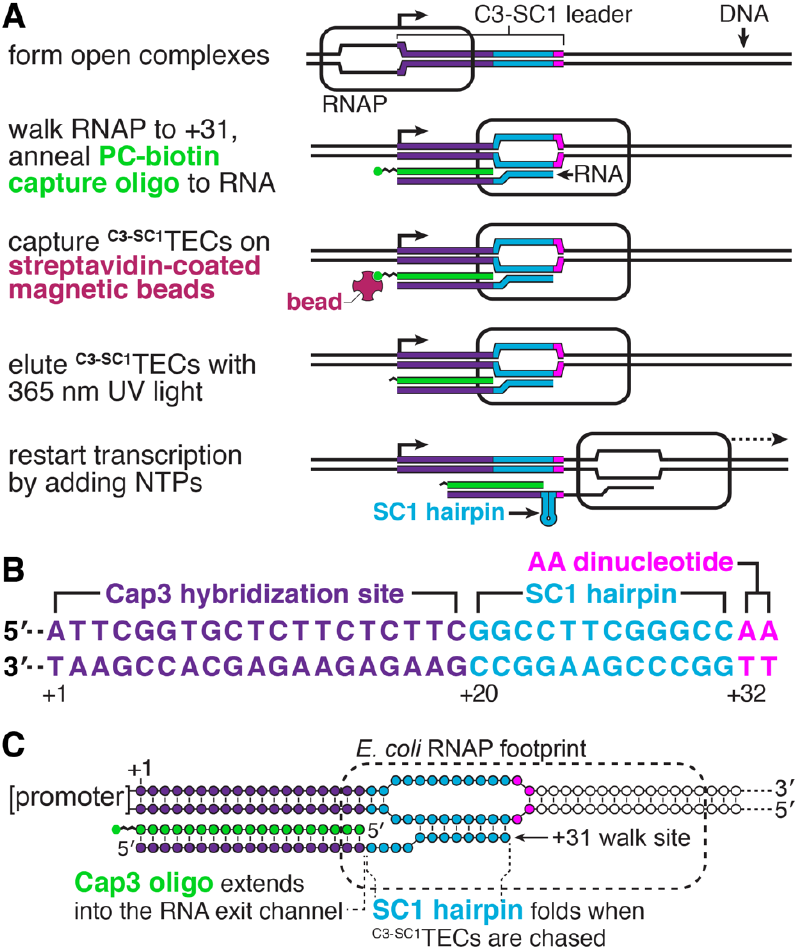
Overview of the ^C3-SC1^TEC purification strategy. *A*, Illustration of the reversible immobilization strategy for isolating ^C3-SC1^TECs. *B*, Annotated sequence of the C3-SC1 leader. *C*, Architecture of ^C3-SC1^TECs. RNAP, RNA polymerase; TEC, transcription elongation complex; PC, photocleavable.

To facilitate this procedure, we designed a 33 nt 5’ leader sequence, called C3-SC1 (<u>C</u>apture sequence <u>3</u>-<u>S</u>tructure <u>C</u>assette <u>1</u>) (Figure 1B). C3-SC1 is composed of a 30 nt A-less cassette embedded between an initiating A_+1_ nucleotide and an A_+32_A_+33_ dinucleotide. Initiating transcription in the presence of an ApU dinucleotide and in the absence of ATP therefore positions RNAP at C_+31_. The C3-SC1 leader contains two functional elements: Nucleotides +1 to +19, called C3 (<u>C</u>apture sequence <u>3</u>), comprise an oligonucleotide hybridization site that can be used to anneal a functionalized oligo to nascent RNA (Figure 1A, B). Nucleotides +20 to +31 comprise a four GC-pair RNA hairpin with a UUCG tetraloop (17,18), referred to here as SC1 (<u>S</u>tructure <u>C</u>assette <u>1</u>), that sequesters this region of the leader when RNAP is chased from the C_+31_ synchronization site (Figure 1A, B). Several considerations were taken into account when designing the C3 sequence because it functions both as an initial transcribed sequence and as an oligo hybridization site. First, occurrences of the YG dinucleotide, which induces transcription pausing (19-23), were limited to locations that do not impact escape from the *λ*P_R_ promoter (24). Second, a G_+5_G_+6_ dinucleotide was included because it favors productive initiation from the *λ*P_R_ promoter (24). Third, homopolymeric sequences were limited to a maximum of two nucleotides to minimize the potential for reiterative transcription initiation (25-28). Fourth, the C3 RNA sequence and its DNA complement are predicted to be unstructured by the ‘fold’ algorithm of the RNAStructure RNA secondary structure prediction software (29). Fifth, the calculated T_m_ of the complementary DNA oligonucleotide (Cap3) is 47.9 *°*C in our transcription conditions to promote efficient hybridization to nascent RNA at 37 *°*C.

In the sections below, TECs that have been synchronized using the C3-SC1 leader and contain the Cap3 oligonucleotide are referred to as ^C3-SC1^TECs (Figure 1C). The spatial organization of nucleic acids within ^C3- SC1^TECs was also considered when designing the C3-SC1 leader. The footprint of RNAP on RNA is ∼14 nts (30). Therefore, when RNAP is positioned at the C_+31_ synchronization site, approximately 17 nts of nascent RNA have emerged from the TEC, assuming RNAP does not backtrack (Figure 1C). The 19 nt-long Cap3 oligonucleotide was designed so that approximately 2 bp of the Cap3-nascent RNA hybrid extends into the RNA exit channel of RNAP. This preserves the register of the RNA 3’ end and RNAP active center because nucleic acid structures at the RNA exit channel prevent RNAP from backtracking (31-33). In addition to functioning as a handle for immobilizing ^C3-SC1^TECs and inhibiting backtracking, the Cap3 oligonucleotide also blocks the C3 sequence from forming base pairs with downstream RNA. Similarly, SC1 forms an RNA hairpin once it emerges from RNAP. In this way, nearly all of the C3-SC1 transcript becomes sequestered in base pairs once RNAP transcribes beyond the synchronization site (Figure 1B).

To be maximally useful, ^C3-SC1^TECs must 1) be homogenous with a 1:1:1 RNAP:DNA:RNA ratio, 2) remain stably associated with DNA during purification, 3) be transcriptionally active, and 4) resume transcription synchronously. In the sections below, we show that ^C3-SC1^TECs isolated using our procedure meet each of these criteria.

### ^C3-SC1^TECs can be immobilized on streptavidin-coated beads

The purification strategy shown in Figure 1 separates ^C3- SC1^TECs from open promoter complexes and free DNA based on the ability of a biotinylated capture oligonucleotide (Cap3) to hybridize to nascent RNA. To assess the amount of Cap3 needed to efficiently form ^C3-SC1^TECs, we generated ^C3- SC1^TECs with variable concentrations Cap3 and determined the fraction of template DNA that could be immobilized on streptavidin-coated magnetic beads. Because Cap3 is the only biotinylated molecule in this experiment, a template DNA molecule can only bind to streptavidin-coated beads if it contains a ^C3-SC1^TEC. For this initial experiment, open promoter complexes were formed using a near-saturating excess of *E. coli* RNAP relative to 10 nM DNA template (34). Consequently, virtually every template DNA molecule contained an open promoter complex and could potentially yield a ^C3-SC1^TEC if RNAP escaped the promoter and the Cap3 oligonucleotide hybridized to the nascent RNA. The fraction of DNA retained on the magnetic beads therefore reflects the approximate upper bound of ^C3-SC1^TEC capture efficiency in our standard *in vitro* transcription conditions. ^C3-SC1^TECs were formed by initiating transcription in the absence of ATP to walk RNAP to the C_+31_ synchronization site (Figure 1). This reaction proceeded at 37 *°*C for 20 minutes to allow Cap3 to bind to nascent RNA before the reaction was mixed with streptavidin-coated beads. Under these conditions, 36-48% of ^C3-SC1^TECs were captured on the beads when biotinylated Cap3 oligo was present at a concentration of 12.5, 25, or 50 nM (Figure 2). In contrast, >97% of ^C3-SC1^TECs remained in the supernatant when a non-biotinylated version of Cap3 was included in the transcription reaction (Figure 2). Given that Cap3 concentration did not meaningfully change the efficiency of ^C3-SC1^TEC capture, all subsequent experiments in this work were performed using 25 nM Cap3 oligonucleotide.

**Figure 2.**
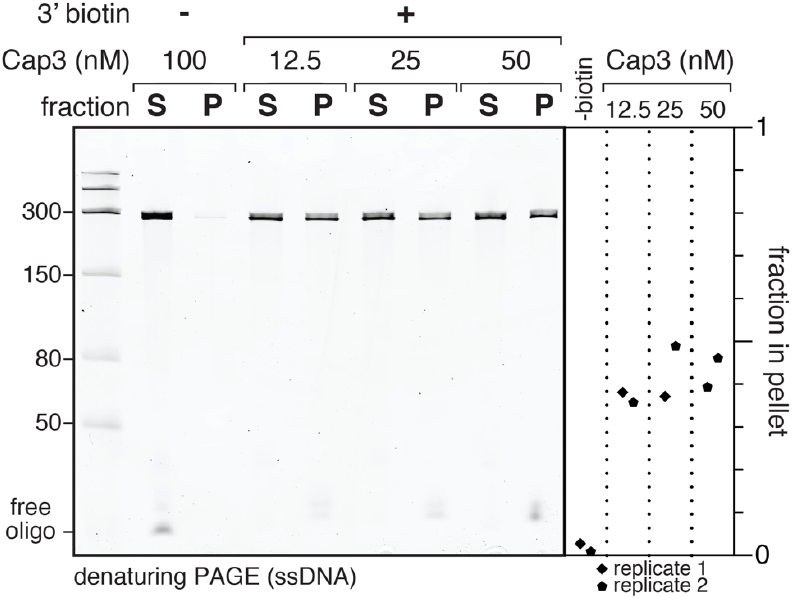
Immobilization of ^C3-SC1^TECs. Denaturing PAGE analysis of DNA from ^C3-SC1^TECs that were assembled using variable amounts of the Cap3_BioTEG or Cap3_NoMod oligonucleotides (Table S1), immobilized on streptavidin-coated magnetic beads, and partitioned into pellet and supernatant fractions. The plot shows the fraction of DNA that was retained on the streptavidin-coated beads. S, supernatant; P, pellet.

### Stringent transcription initiation conditions enrich for uniform ^C3-SC1^TECs

The experiments in Figure 2 used a near-saturating concentration of RNAP to assess the efficiency limit of ^C3- SC1^TEC immobilization. However, under these conditions the exact composition of the immobilized complexes is poorly defined because each DNA molecule can contain both a ^C3-SC1^TEC and an open promoter complex. Ideally, each DNA molecule should contain a maximum of one RNAP so that the RNAP:DNA:RNA ratio is 1:1:1. After evaluating several variations of the ^C3-SC1^TEC purification procedure, we identified stringent transcription initiation conditions that yield >95% pure ^C3-SC1^TECs with a 1:1:1 RNAP:DNA:RNA ratio (Figures 3 and S1, see Experimental Procedures for complete details): First, we used a sub-saturating concentration of RNAP when forming open promoter complexes to minimize non-specific DNA binding. We previously used this strategy when purifying arrested TECs (34). Next, we challenged the open promoter complexes with 20 μg/ml heparin to sequester free RNAP and RNAP that was weakly associated with DNA. Finally, we initiated single-round transcription by simultaneously adding MgCl_2_ and the antibiotic rifampicin to the reaction. This strategy limits the occurrence of abortive cycling during the Cap3 oligonucleotide hybridization and ^C3-SC1^TEC immobilization steps. After assembling and immobilizing ^C3-SC1^TECs using these conditions, the immobilized complexes were washed to remove excess transcription reaction components and eluted by irradiating the sample with 365 nm UV LEDs (∼10 mW/cm^2^) for 5 minutes (Figure 3A).

**Figure 3.**
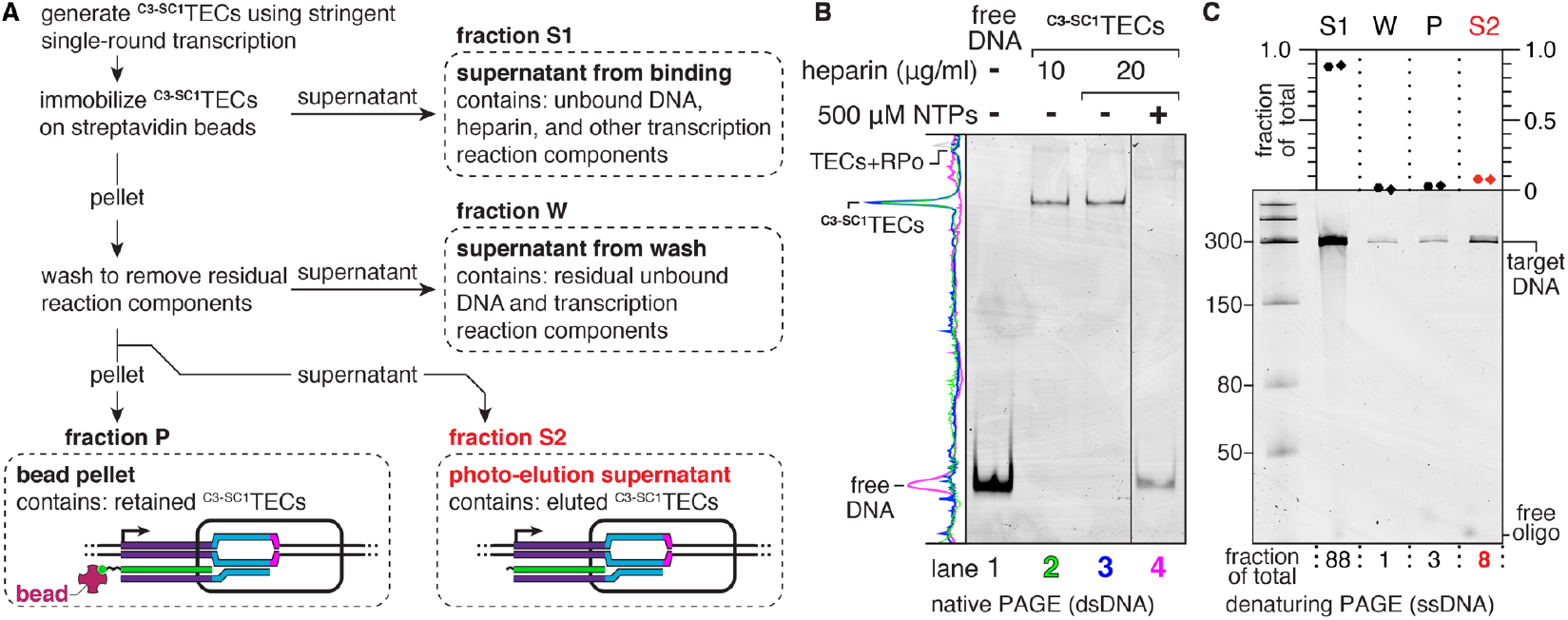
Purification of homogenous ^C3-SC1^TECs by reversible immobilization. *A*, Workflow of the ^C3-SC1^TEC purification procedure. The source of each fraction (S1, W, P, S2) that was collected for the experiment in panel *C* is indicated. *B, EMSA* of ^C3-SC1^TECs that were purified using transcription initiation conditions that favor a 1:1:1 RNAP:DNA:RNA ratio, before and after the resulting complexes were chased by adding NTPs. Lanes 2 and 3 show ^C3-SC1^TECs that were prepared using two concentrations of heparin; three additional independent ^C3-SC1^TEC preparations are shown in Figure S1A. The solid vertical line between lanes 3 and 4 indicates a gel splice to remove lanes that were left empty to avoid the potential for NTP contamination in the -NTP samples. *C*, Denaturing PAGE analysis of the ^C3-SC1^TEC purification procedure from *A*. The plot shows the fraction of total DNA that was recovered in fractions S1, W, P, and S2. TEC, transcription elongation complex; RPo, open promoter complex; S1, supernatant 1; W, wash; P, pellet; S2, supernatant 2.

^C3-SC1^TECs that were prepared using this stringent transcription initiation procedure contained a fast-migrating major band (>95% of complexes) and a slow-migrating minor band (<5% of complexes) when assessed by EMSA (Figure 3B). We previously observed a similar migration pattern when assessing the purity of arrested TECs (34); the fast-migrating band corresponded to pure TECs and the slow-migrating band corresponded to TECs with an associated open promoter complex. To test whether this interpretation applied to ^C3-SC1^TECs, we prepared ^C3-SC1^TECs using a saturating concentration of RNAP and omitted heparin; these conditions favor the formation of new open complexes after RNAP escapes the promoter. As expected, ^C3-SC1^TECs prepared using these conditions contained ∼60% slow-migrating complexes and ∼40% fast-migrating complexes (Figure S1B). We therefore conclude that the fast-migrating band corresponds to pure ^C3-SC1^TECs and the slow-migrating corresponds to ^C3-SC1^TECs with an associated open promoter complex. ^C3-SC1^TECs prepared using the stringent transcription initiation conditions described above are therefore >95% homogenous. Importantly, incubating ^C3- SC1^TECs with 500 μM NTPs caused the template DNA to migrate with the same mobility as free DNA, which indicates that RNAP was released from DNA and that the resulting complexes are active (Figure 3B).

The upper bound for ^C3-SC1^TEC yield is 40-50% when a near-saturating amount of RNAP is used for transcription (Figure 2). We anticipated that the stringent conditions required to isolate homogenous complexes would reduce the yield of ^C3-SC1^TECs. To test this, we evaluated the distribution of input DNA across four purification fractions (Figure 3A). As expected, ^C3-SC1^TECs that were purified using the stringent transcription initiation conditions described above contained only 8% of the total input DNA (Figure 3C). Sequestering free RNAP with a competitor DNA template instead of heparin increased the yield to 16% but reduced the fraction of ^C3-SC1^TECs with a 1:1:1 RNAP:DNA:RNA ratio from >95% to ∼88% (Figure S1, B, C, and D). Omitting rifampicin had virtually no effect on yield (Figure S1E). The low yield of the optimized ^C3-SC1^TEC preparation is therefore primarily due to the use of a sub-saturating RNAP concentration and the use of heparin to sequester excess and weakly bound RNAP. However, this limitation is offset by the high purity and robust activity of the resulting complexes.

### ^C3-SC1^TECs resume transcription synchronously

The observation that incubating purified ^C3-SC1^TECs with NTPs releases RNAP from DNA suggests that the complexes are transcriptionally active. To assess the nucleic acid composition and reactivation kinetics of ^C3-SC1^TECs more precisely, we prepared ^C3-SC1^TECs that contained ^32^P-labeled RNA and performed a transcription time course. In this experiment, the template DNA contained an etheno-dA transcription stall site 28 nt downstream of the synchronization site to halt transcription uniformly (35,36). Before NTPs were added, virtually all ^C3-SC1^TECs contained either a 31 or 32 nt-long RNA (Figure 4A). The presence of a single discrete band at the etheno-dA stall site indicates that the 32 nt RNA is due to transcription 1 nt beyond the C_+31_ synchronization site, likely due to either trace ATP or misincorporation, rather than transcription start site variability (Figure 4A). When transcription was reactivated by adding NTPs to 500 μM, 94% of ^C3-SC1^TECs transcribed to the etheno-dA stall site in <15 seconds (Figure 4B). Virtually all ^C3-SC1^TECs that did not transcribe to the etheno-dA stall site within 15 seconds contained a 32 nt-long RNA (Figure 4B). Although this sub-population resumed transcription slowly, >98% of ^C3-SC1^TECs transcribed to the etheno-dA stall site within one minute. Given that purified ^C3-SC1^TECs do not contain free DNA and are >98% active, *in vitro* transcription reactions that use these complexes maintain a quantitative relationship between the template DNA and RNA transcript.

**Figure 4.**
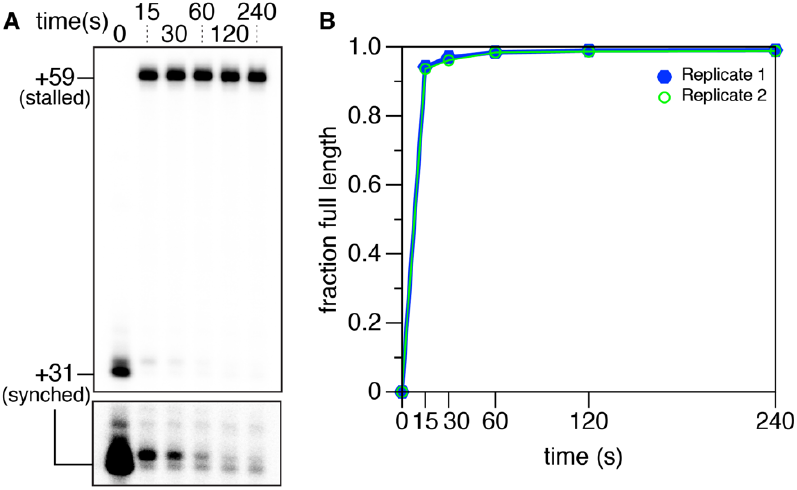
Analysis of ^C3-SC1^TEC reactivation kinetics. A, Transcription time course to assess the activity of ^C3- SC1^TECs. RNAP stalls at +59 due to the presence of an etheno-dA modification in the template DNA. The lower cut-out is adjusted to show trace +31 and +32 RNAs in the 15 s to 240 s time points; the entire gel is shown with this darker grayscale setting in Figure S2A. B, Plot showing the fraction of ^C3-SC1^TECs that transcribed to the +59 stall site at each time point.

### ^C3-SC1^TECs are compatible with solid-phase transcription reactions

Our analysis of ^C3-SC1^TECs that had been eluted from streptavidin-coated beads established their purity and activity. However, for some applications it is desirable to perform *in vitro* transcription as a solid-phase reaction. To assess whether ^C3-SC1^TECs are compatible with solid-phase transcription, we prepared ‘permanently’ immobilized ^C3-SC1^TECs using a biotinylated Cap3 oligonucleotide that did not contain a photocleavable spacer modification (Cap3_BioTEG, Table S1) and a DNA template that contained an etheno-dA stall site. >99% of ^C3- SC1^TECs remained immobilized both before and after the addition of NTPs, and >99% of ^C3-SC1^TECs transcribed to the etheno-dA stall site (Figure 5). We therefore conclude that ^C3-SC1^TECs are fully compatible with solid-phase transcription.

**Figure 5.**
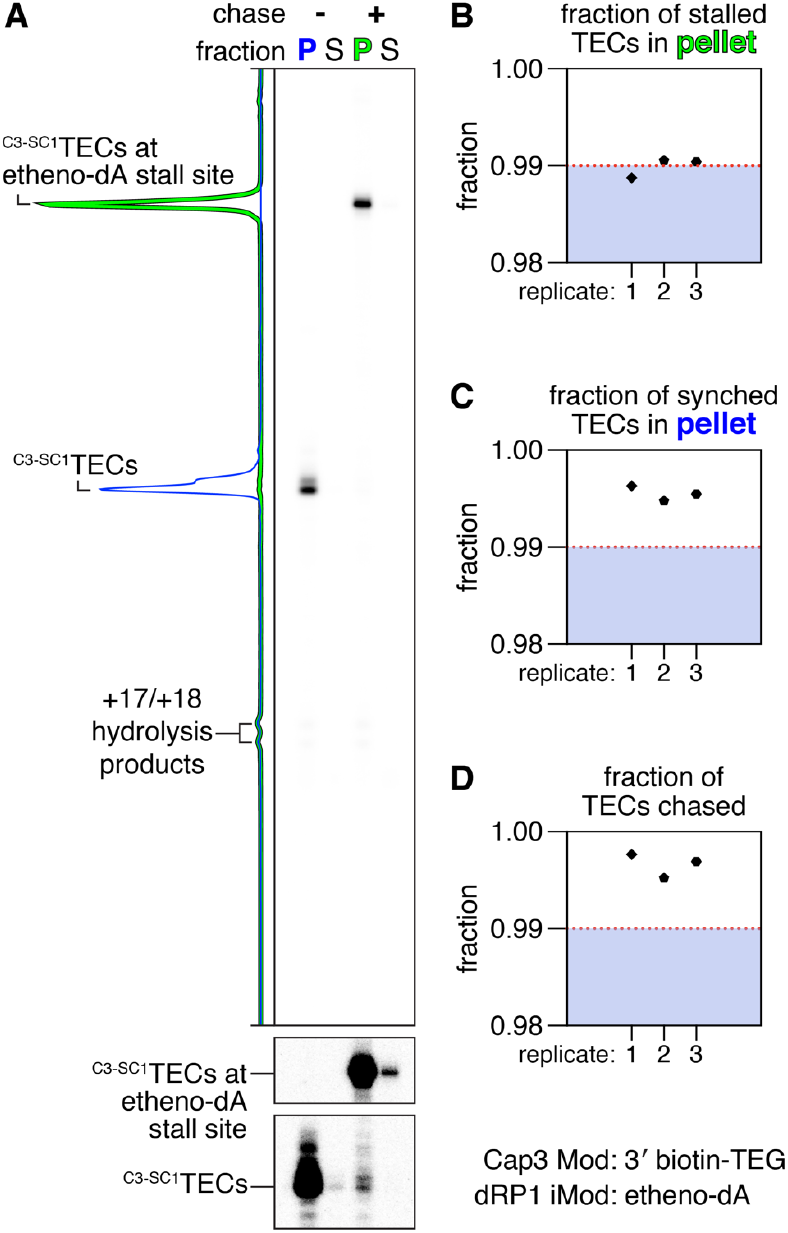
Solid-phase-transcription using ^C3-SC1^TECs. *A*, ^C3-SC1^TECs containing the Cap3_BioTEG oligonucleotide were immobilized on streptavidin-coated magnetic beads and partitioned into pellet and supernatant fractions before and after NTPs were added to reactivate transcription. RNAP stalls at +59 and is retained on DNA due to the presence of an etheno-dA modification in the template DNA. The lower cut-out is adjusted to show trace synchronized and etheno-dA stalled RNAs in the supernatant fractions; the entire gel is shown with this darker grayscale setting in Figure S2B. The fraction of stalled (*B*) and synchronized (C) ^C3-SC1^TECs that were retained in the pellet are shown. *D*, Plot showing the fraction of ^C3-SC1^TECs that transcribed to the +59 stall site after NTPs were added.

We detected trace amounts of transcriptionally inactive 15 to 18 nt-long RNAs (primarily 17 and 18 nts) in both this assay and the transcription time course assay (Figure S2). With the exception of the 15 nt RNA, these transcripts were retained on the beads and are therefore annealed to the Cap3 oligonucleotide. We therefore conclude that these short transcripts are hydrolysis products of the C3 region of the C3-SC1 transcript that occur during the purification of ^C3-SC1^TECs.

### The C3-SC1 leader is compatible with riboswitch functional assays

When RNAP transcribes beyond the C_+31_ synchronization site, 31 of 33 nucleotides in the C3-SC1 transcript are sequestered in base pairs with the Cap3 oligonucleotide or in a fast-folding RNA hairpin (Figure 1). The purpose of this design is to limit interactions between the C3-SC1 transcript and downstream RNA structures that could potentially interfere with RNA function. To determine whether this strategy was successful, we assessed how the C3-SC1 transcript affects the transcription antitermination activity of the *Clostridium beijerinckii pfl* ZTP riboswitch (37) and the *Clostridiales bacterium* (*Cba*) ppGpp riboswitch (38). The C3-SC1 leader was appended to the 5’ end of each riboswitch sequence (Figure 6, A and D). Transcription using ^C3-SC1^TECs reduced terminator readthrough from 19% to 9% for the *pfl* riboswitch and from 38% to 7% for the *Cba* riboswitch (Figure 6, B and E). The C3-SC1 transcript did not meaningfully reduce the amount of transcription antitermination observed for the *pfl* ZTP riboswitch at the highest ZMP concentration tested (1 mM ZMP), and consequently increased fold-change from 3.6 to 7.1 over this concentration range (Figure 6, B and C). In contrast, the C3-SC1 transcript decreased the amount of transcription terminator readthrough observed for the *Cba* ppGpp riboswitch at all concentrations tested (Figure 6, E and F). Although the difference between terminator readthrough without ppGpp and at saturating ppGpp remained approximately constant, (37% for ^C3-SC1^TECs, 32% for leader-less riboswitches), the reduced amount of ligand-independent terminator readthrough observed for ^C3-SC1^TECs increased fold-change from 1.9 to 6.0. Given these findings, we conclude that transcription using ^C3-SC1^TECs is generally perturbative to RNA function, but that these perturbations do not necessarily interfere with function.

**Figure 6.**
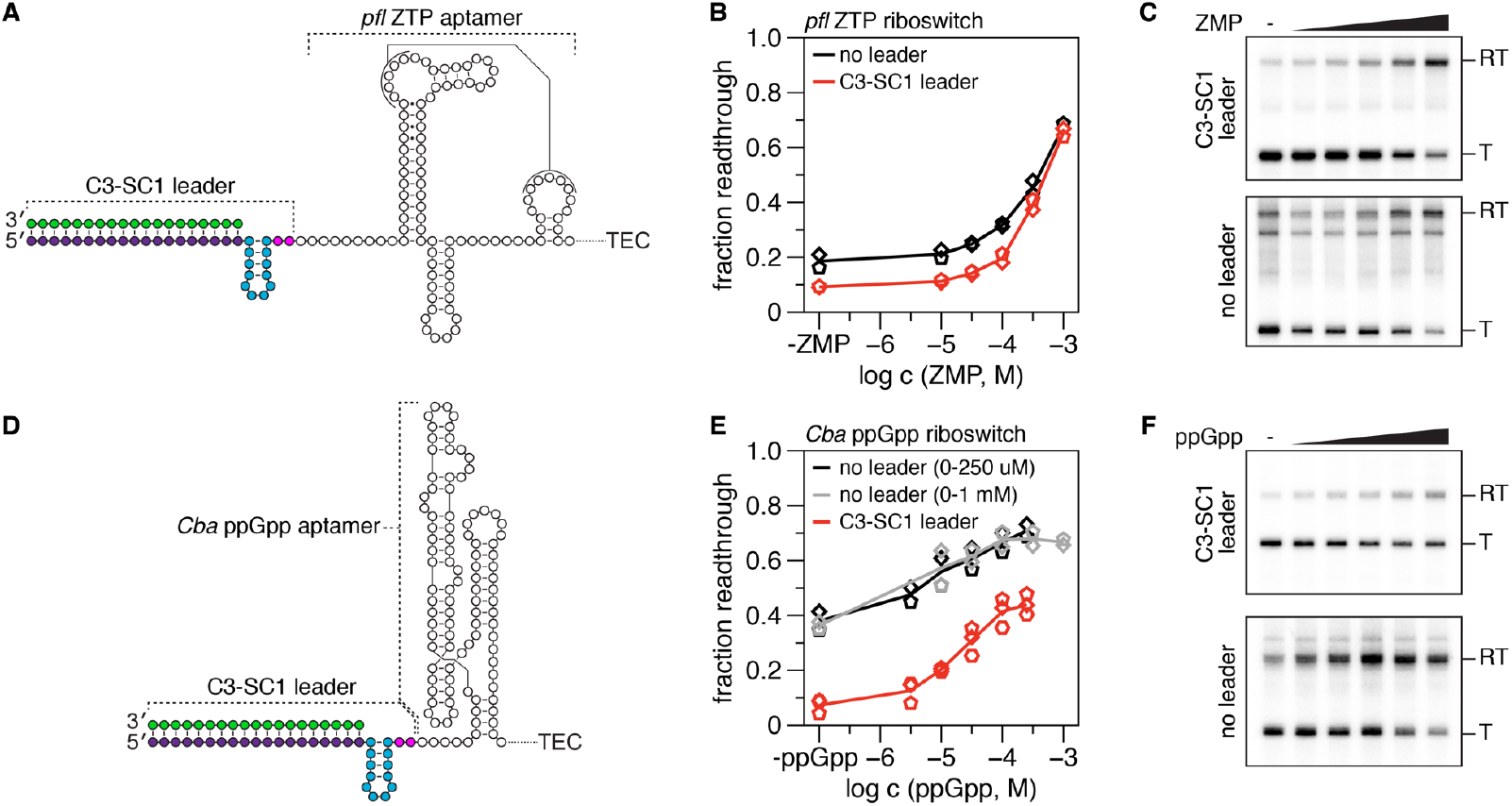
Transcriptional riboswitch assays using ^C3-SC1^TECs. A, Illustration of the Clostridium beijerinckii pfl ZTP aptamer with the C3-SC1 transcript appended to its 5’ end. B, ZMP dose-response curves for the pfl ZTP riboswitch with and without the C3-SC1 leader. C, Representative gels for the analysis in B. D, Illustration of the Clostridiales bacterium ppGpp aptamer with the C3-SC1 transcript appended to its 5’ end. E, ppGpp dose-response curves for the Cba ppGpp riboswitch with and without the C3-SC1 leader. F, Representative gels for the analysis in E. 500 μM NTPs were used for all riboswitch assays. T, terminated; RT, readthrough.

## Discussion

We have developed a procedure for isolating synchronized TECs from an *in vitro* transcription reaction using a photoreversible immobilization strategy. Our approach uses a custom 5’ leader sequence called C3-SC1 to generate synchronized TECs that can be immobilized, washed, and recovered by 365 nm UV irradiation (Figure 1). TECs isolated using this procedure (^C3-SC1^TECs) are >95% pure, >98% active, compatible with solid-phase transcription, and can be used in biochemical assays for RNA function. Importantly, the removal of free template DNA molecules during ^C3-SC1^TEC purification establishes a 1:1:1 ratio between RNAP, template DNA, and the nascent RNA transcript, and the high activity of the resulting complexes maintains this ratio when transcription is resumed. These properties make ^C3-SC1^TECs a potentially powerful tool for developing biochemical and biophysical assays that require high-performance, quantitative bacterial *in vitro* transcription.

Our procedure for isolating ^C3-SC1^TECs is optimized for purity and activity, but has three primary limitations: First, the stringent transcription initiation conditions that are used to maximize the purity of ^C3-SC1^TECs reduced the yield to ∼8% of total input DNA. Challenging open promoter complexes with a competitor DNA template instead of heparin doubled the yield but reduced the purity of the ^C3-SC1^TECs from >95% to ∼88% and is therefore not ideal. Second, excess Cap3 oligonucleotide cannot be removed from the ^C3-SC1^TEC preparation easily because it contains the attachment chemistry needed to pull down the target complexes. The presence of free Cap3 oligonucleotide in ^C3-SC1^TEC preparations did not interfere with the transcription termination and antitermination functions of the riboswitches that were assessed in this work (Figure 6). However, the effect of free Cap3 oligonucleotide on downstream biochemical assays will be application-dependent. Third, the procedure that we have described is not necessarily compatible with all bacterial promoters. In the experiments described above we used a custom σ^70^ promoter that is primarily derived from the λP_R_ promoter (34), which generates long-lived, heparin-resistant open complexes (39). Promoters that generate short-lived, heparin-sensitive open complexes will not be compatible with our procedure. Given these limitations, ^C3-SC1^TECs will be most useful for experiments that demand the use of pure, synchronous TECs to transcribe complex RNA molecules, but which do not require the use of a specific promoter or obtaining concentrated transcription complexes. In this way, the procedure we have described complements existing scaffold-based strategies for assembling high-purity synchronized TECs from DNA and RNA oligonucleotides (40,41). Nucleic acid scaffolds can be used prepare high-purity synchronized TECs at micromolar concentrations, but can only be used to synthesize long transcripts if the scaffold DNA is ligated to a second template DNA molecule (41). In contrast, ^C3-SC1^TECs can be assembled on long or complex template DNA molecules because they are generated by promoter-directed transcription initiation using standardized sequences.

Because the ability of leader transcripts to perturb RNA function is well-established (42), the C3-SC1 leader was designed to minimize base-pairing interactions with downstream RNA. Nonetheless, the C3-SC1 transcript perturbed the transcription antitermination activity of ZTP-and ppGpp-sensing transcriptional riboswitches (Figure 6). Although this outcome was not surprising, its nature was unexpected: transcription using ^C3-SC1^TECs reduced the probability that RNAP bypasses the transcription termination site in the absence of ligand but did not interfere with the ability of either riboswitch to antiterminate transcription in the presence of ligand. In effect, ^C3-SC1^TECs improved the biochemical activity of both riboswitch systems by increasing the fold-change of the switches. One possible explanation for this effect is that the C3-SC1 transcript destabilizes the apo ZTP and ppGpp aptamers so that terminator hairpin folding is more efficient, but does not cause the aptamers to catastrophically misfold. Regardless of the mechanistic origin of this effect, its occurrence underscores the importance of evaluating how the C3-SC1 transcript impacts RNA function when using ^C3-SC1^TECs. Given that the C3-SC1 transcript did not interfere with the function of either of the structurally distinct riboswitches assessed in this work, it is likely that ^C3-SC1^TECs will be compatible with many RNA targets. Furthermore, ^C3-SC1^TECs may be useful for manipulating the activity of riboswitch RNAs *in vitro* if the functional enhancements that we observed for the ZTP and ppGpp riboswitches are generalizable to other systems.

## Experimental Procedures

### Oligonucleotides

All oligonucleotides were purchased from Integrated DNA Technologies. A detailed description of all oligonucleotides including sequence, modifications, and purifications is presented in Table S1. Out of an abundance of caution, oligonucleotides that contained a PC spacer modification were handled under low intensity 592 nm light from SimpleColor Amber LEDs (Waveform Lighting) set to 40% intensity using a FilmGrade Flicker-Free LED Dimmer (Waveform Lighting) and stored as single-use aliquots.

### Proteins

Vent (exo-) DNA polymerase, Q5 High-Fidelity DNA Polymerase, *Sulfolobus* DNA Polymerase IV, and *E. coli* RNA Polymerase holoenzyme were purchased from New England Biolabs. NusA was a gift from J. Roberts (Cornell University).

### DNA template sequences

DNA template sequences are available in Table S2. Fully annotated versions of DNA templates 1 (https://benchling.com/s/seq-FUAWrFQwiHR3aJmAUExl), 2 (https://benchling.com/s/seq-3hXcRKbqeVo9u4Hhl2oI), 3 (https://benchling.com/s/seq-wIQ0VHiDjt1ahpC9ELjD), 4 (https://benchling.com/s/seq-XFA473UmWVvfLaubWzXa), 5 (https://benchling.com/s/seq-2QwZ9VceV9l3uarzKCKT) and 6 (https://benchling.com/s/seq-LI54uN63QQ9QyAAJXIAi) are available at Benchling.

### DNA template preparation

Table S3 provides details on the oligonucleotides and processing steps used for every DNA template preparation used in this work. Unmodified DNA templates were PCR amplified from plasmid DNA using Vent (exo-) DNA polymerase and purified by agarose gel extraction exactly as described previously (34). DNA templates that contained an internal etheno-dA transcription stall site were PCR amplified from an unmodified linear dsDNA using Q5 High-Fidelity DNA Polymerase; the unmodified linear dsDNA that was used as a template for these reactions was PCR amplified from plasmid DNA using Vent (exo-) DNA polymerase and purified by agarose gel extraction. After PCR amplification, DNA containing an internal etheno-dA modification was purified using a QIAquick PCR Purification Kit (Qiagen), processed by translesion DNA synthesis using a mixture of *Sulfolobus* DNA Polymerase IV and Vent (exo-) DNA polymerase, and purified a second time using a QIAquick PCR Purification Kit; these steps were performed exactly as described previously (34). A step-by-step protocol describing this procedure is available (43).

### Preparation of streptavidin-coated magnetic beads

5 μl of 10 mg/ml Dynabeads MyOne Streptavidin C1 beads (Invitrogen) per 25 μl sample volume were prepared in bulk exactly as described previously (34). The resulting beads were resuspended at a concentration of ∼2 μg/μl in Buffer TX (1X Transcription Buffer (20 mM Tris-HCl (pH 8.0), 50 mM KCl, 1 mM DTT, 0.1 mM EDTA (pH 8.0)) supplemented with 0.1% Triton X-100), split into in 25 μl aliquots, and stored on ice until use.

### Capture oligo dose assays

For the capture oligo dose assay in Figure 2, 25 μl *in vitro* transcription reactions containing 1X Transcription Buffer, 0.1 mg/ml BSA, 10 mM MgCl_2_, 100 μM ApU dinucleotide (Dharmacon), 1X UGC Start NTPs (25 μM UTP, 25 μM GTP, 25 μM CTP, all NTP mixtures were prepared using High Purity rNTP solutions (Cytiva)), 10 nM DNA Template 1 (Tables S2 and S3), 0.024 U/μl *E. coli* RNAP holoenzyme, and Cap3 oligonucleotide (either 12.5, 25, or 50 nM Cap3_BioTEG or 100 μM Cap3_NoMod, Table S1) were prepared in a 1.7 ml microcentrifuge tube on ice; at this point the total reaction volume was 22.5 μl due to the omission of 10X (100 μg/ml) rifampicin.

Transcription reactions were placed in a dry bath set to 37 *°*C for 20 minutes to walk RNAP to C_+31_ of the C3-SC1 leader. After 20 minutes, rifampicin was added to a final concentration of 10 μg/ml and the sample was incubated at 37 *°*C for 5 additional minutes to block new transcription initiation. Pre-equilibrated, room-temperature streptavidin-coated magnetic beads were placed on a magnetic stand and storage buffer was removed. The streptavidin-coated magnetic beads were resuspended with the transcription reaction by pipetting and incubated at room temperature with rotation for 15 minutes. After 15 minutes, the bead binding mixture was spun briefly in a Labnet Prism mini centrifuge (Labnet International) by quickly flicking the switch on and off so that liquid was removed from the tube cap. The sample was placed on a magnetic stand for 1 minute, the supernatant was transferred to a 1.7 ml microcentrifuge tube containing 125 μl Stop Solution (0.6 M Tris-HCl (pH 8.0), 12 mM EDTA (pH 8.0)), and the pellet was resuspended in 25 μl of 95% deionized formamide and 10 mM EDTA, heated at 95 *°*C for 5 minutes, placed on a magnet stand for 1 min, and the supernatant was collected and mixed with 125 μl of Stop Solution. The samples were then processed by phenol:chloroform extraction and ethanol precipitation, and fractionated by denaturing Urea-PAGE as described below in the section *Collection, processing, and denaturing PAGE of* ^*C3-SC1*^*TEC purification fractions*.

### Preparation of ^C3-SC1^TECs

The transcription conditions used for the preparation of ^C3-SC1^TECs were selected to enrich for active transcription complexes. First, a sub-saturating amount of *E. coli* RNAP (determined previously (34)) was used to minimize non-specific DNA binding during open complex formation. Second, open promoter complexes were challenged with heparin (Sigma-Aldrich, catalog # H5515) to sequester free RNAP and enrich for heparin-resistant open complexes. Third, transcription was limited to a single-round using rifampicin so that transcription was not actively occurring during the Cap3 oligonucleotide hybridization and streptavidin-coated bead binding steps. Below, the final validated procedure is detailed first and other variations of the procedure that were performed during protocol development are then described with reference to the figures in which they were used.

25 μl *in vitro* transcription reactions containing 1X Transcription Buffer, 0.1 mg/ml BSA, 100 μM ApU dinucleotide, 1X UGC Start NTPs (25 μM UTP, 25 μM GTP, 25 μM CTP), 10 nM template DNA, 0.016 U/μl *E. coli* RNAP holoenzyme, and 25 nM Cap3_PCBioTEG oligonucleotide (Table S1) were prepared in a 1.7 ml microcentrifuge tube on ice; at this point the total reaction volume was 20 μl due to the omission of 10X (200 μg/ml) heparin and 10X Start Solution (100 mM MgCl_2_, 100 μg/ml rifampicin). The Cap3_PCBioTEG oligonucleotide was added to the reaction last under 592 nm amber light, and all sample handling until the 365 nm UV irradiation step was performed under 592 nm amber light. When preparing radiolabeled ^C3-SC1^TECs, 0.2 μCi/μl [σ-^32^P]UTP (PerkinElmer) was included in the transcription reaction.

Transcription reactions were placed in a dry bath set to 37 *°*C for 20 minutes to form open promoter complexes. After 20 minutes, 2.5 μl of 200 μg/ml heparin per sample volume was added to the reaction, and the sample was mixed by pipetting and incubated at 37 *°*C for 5 minutes to sequester free RNAP and enrich for heparin-resistant open promoter complexes; the final concentration of heparin was 20 μg/ml. After 5 minutes, 2.5 μl of room temperature 10X Start Solution per sample volume was added to the transcription reaction for a final concentration of 10 mM MgCl_2_ and 10 μg/ml rifampicin. The transcription reaction was mixed by pipetting and incubated at 37 *°*C for 20 minutes to walk RNAP to the C_+31_ synchronization site and hybridize the Cap3 oligonucleotide. At this time, tubes containing 25 μl of ∼2 μg/μl pre-equilibrated streptavidin-coated magnetic beads and Buffer TMW (1X Transcription Buffer, 10 mM MgCl_2_, and 0.05% Tween-20) were placed at room temperature.

After ∼18 minutes, the magnetic beads were placed on a magnetic stand and storage buffer was removed. After 20 minutes, the beads were resuspended with the transcription reaction by pipetting and incubated in the dark at room temperature with rotation for 15 minutes. After 15 minutes, the bead binding mixture was spun briefly in a Labnet Prism mini centrifuge by quickly flicking the switch on and off so that liquid was removed from the tube cap, but the speed of the mini centrifuge remained as low as possible. The sample was placed on a magnet stand for 1 minute and the supernatant was removed; this supernatant, which contained any transcription reaction components that did not bind the beads, is referred to as fraction S1. The 1.7 ml tube containing the sample was removed from the magnet stand and the beads were gently resuspended in either 250 μl (for single samples) or 1 ml (for bulk samples) of room temperature Buffer TMW and incubated in the dark at room temperature with rotation for 5 minutes. The sample was placed on a magnet stand for 1 minute and the supernatant was removed; this supernatant, which contains residual transcription reaction components, is referred to as fraction W. In all experiments except the riboswitch functional assays described below, the beads were gently resuspended in 25 μl of Buffer TMW per sample volume by pipetting so that the bead concentration was ∼2 μg/μl, placed in a custom-built 365 nm UV LED irradiator for 1.7 ml microcentrifuge tubes, and exposed to ∼10 mW/cm^2^ 365 nm UV light from four directions for 5 minutes (34); all sample handling was performed under normal light after this step was complete. After UV irradiation, the sample was returned to the magnet stand for 1 minute and the supernatant was collected. The bead pellet, which contains any ^C3-SC1^TECs that were not eluted by 365 nm UV irradiation, is referred to as fraction P. The collected supernatant, which contains eluted ^C3- SC1^TECs, is referred to as fraction S2.

Several variations of this protocol were performed. In addition to preparing ^C3-SC1^TECs with 20 μg/ml heparin, ^C3- SC1^TECs were also prepared with 10 μg/ml and 15 μg/ml heparin without any detectable difference (Figure 3, Figure S1A). In preliminary experiments, open promoter complexes were incubated with 15, 22.5, or 30 nM competitor DNA containing an etheno-dA lesion for 20 minutes in place of heparin (Figure S1, B, C, and D). In control experiments ^C3-SC1^TECs were prepare using 5 nM DNA template, 0.024 U/μl *E. coli* RNAP holoenzyme, and no heparin or competitor DNA (Figure S1B) and without rifampicin (FigureS1E).

### Analysis of ^C3-SC1^TECs by EMSA

EMSAs were performed exactly as described previously (34) by loading samples onto a 0.5X TBE 5% polyacrylamide gel prepared for a Mini-PROTEAN Tetra Vertical Electrophoresis Cell (Bio-Rad) using ProtoGel acrylamide (National Diagnostics). To assess purified ^C3-SC1^TECs, 15 μl of a 25 μl sample was mixed with 3 μl of 6X Native DNA Loading Dye (30% (v/v) glycerol, 10 mM Tris-HCl (pH 8.0), 0.01% (w/v) bromophenol blue) and loaded onto the gel. To assess ^C3-SC1^TECs after reactivating transcription, a 25 μl sample was mixed with 0.5 μl of 500 μg/ml rifampicin, incubated at 37 *°*C for 2 minutes, mixed with 0.5 μl of 25 mM NTPs, and incubated at 37*°* C for 2 minutes; 15.6 μl of the 26 μl reaction was mixed with 3 μl of 6X Native DNA Loading Dye and loaded onto the gel. After the gel had run for ∼2 h and 45 min at 45 V, the gel was transferred to a plastic dish containing 1X SYBR Gold (Invitrogen) in 0.5X TBE, stained for 10 minutes with rocking, and scanned on a Sapphire Biomolecular imager using the 488 nm/518BP22 setting.

### Collection, processing, and denaturing PAGE of ^C3-SC1^TEC purification fractions

^C3-SC1^TEC purification fractions were collected and processed as follows: Fraction S1 was mixed with 125 μl of Stop Solution. Fraction W was mixed with 5 μl of 0.5 M EDTA (pH 8.0). To recover immobilized nucleic acids from Fraction P, the bead pellet was resuspended in 25 μl of 95% deionized formamide and 10 mM EDTA, heated at 95 *°*C for 5 minutes, placed on a magnet stand for 1 min, and the supernatant was collected and mixed with 125 μl of Stop Solution. Fraction S2 was mixed with 125 μl of Stop Solution. The fractions were extracted by adding an equal volume of Tris (pH 8) buffered phenol:chloroform:isoamyl alcohol (25:24:1, v/v), mixing by vortexing and inversion, and centrifuging at 18,500 × g and 4 *°*C for 5 minutes. The aqueous phase was collected and transferred to a new tube. Nucleic acids were precipitated by adding 0.1 sample volumes of 3 M sodium acetate (pH 5.5), 3 sample volumes of 100% ethanol, and 1 μl of GlycoBlue Coprecipitant (Invitrogen), and chilling at -70 *°*C for 30 minutes. The samples were centrifuged at 18,500 x g and 4 *°*C for 30 minutes, the supernatant was removed, the samples were centrifuged again briefly to pull down residual ethanol, and residual ethanol was removed. The pellet was dissolved in 16 μl of Formamide Loading Dye (90% (v/v) deionized formamide, 1X Transcription Buffer, 0.01% (w/v) bromophenol blue), heated at 95 *°*C for 5 minutes, and snap-cooled on ice for 2 minutes. The samples were then assessed by Urea-Page using an 8% gel prepared with the SequaGel UreaGel 19:1 Denaturing Gel System (National Diagnostics) for a Mini-PROTEAN Tetra Vertical Electrophoresis Cell exactly as described previously (34).

### Transcription kinetics assay

Seven sample volumes of ^C3-SC1^TECs containing [σ-^32^P]UTP-labeled RNA were prepared in bulk using DNA template 2 (Tables S2 and S3) as described in the section *Preparation of* ^*C3-SC1*^*TECs*. The resulting 175 μl pooled sample was mixed with 3.5 μl of 500 μg/ml rifampicin and incubated at 37 *°*C for 2 minutes. To take a zero time point, 25 μl of the pooled sample was removed, mixed with 125 μl of Stop Solution, and kept on ice. The remaining 150 μl pooled sample was mixed with 3 μl of 25 mM NTPs for a final NTP concentration of ∼500 μM NTPs and incubated at 37 *°*C. At each time point (15 s, 30 s, 1 min, 2 min, and 4 min) a 25 μl sample volume was removed from the pooled sample, mixed with 125 μl of Stop Solution, and transferred to ice. The samples were processed as described below in the section *Purification, sequencing gel electrophoresis, and detection of radiolabeled RNA*.

### Immobilized ^C3-SC1^TEC activity assay

Immobilized ^C3-SC1^TECs containing [σ-^32^P]UTP-labeled RNA were prepared in bulk using DNA Template 2 (Tables S2 and S3) as described in the section *Preparation of* ^*C3-SC1*^*TECs* except the Cap3_BioTEG oligonucleotide (Table S1) was used instead of Cap3_PCBioTEG and the photo-elution step was omitted. 50 μl of immobilized ^C3-SC1^TECs were mixed with 1 μl 500 μg/ml rifampicin and incubated at 37 *°*C for 2 minutes. 25 μl of the sample was transferred to a separate 1.7 ml microcentrifuge tube and placed on a magnetic stand for 1 min. The supernatant was transferred to 125 μl of Stop Solution and the beads were resuspended in 25 μl of 95% deionized formamide and 10 mM EDTA. The remaining 25 μl of the sample was mixed with 0.5 μl of 25 mM NTPs for a final concentration of ∼500 μM NTPs, incubated at 37 *°*C for 1 min, and placed on a magnetic stand for 1 min. The supernatant was transferred to 125 μl of Stop Solution and the beads were resuspended in 25 μl of 95% deionized formamide and 10 mM EDTA. To recover immobilized nucleic acids, the pellet fractions were heated at 95 *°*C for 5 minutes, placed on a magnet stand for 1 min, and the supernatant was collected and mixed with 125 μl of Stop Solution. The samples were processed as described below in the section *Purification, sequencing gel electrophoresis, and detection of radiolabeled RNA*.

### Transcription termination assays

Transcription termination assays using ^C3-SC1^TECs were performed as follows: ^C3-SC1^TECs containing [σ-^32^P]UTP-labeled RNA were prepared in bulk using either DNA template 1 or 4 (Tables S2 and S3) as described above in the section *Preparation of* ^*C3-SC1*^*TECs*, except that ^C3-SC1^TECs were eluted into either 19.5 μl per sample volume of Elution Buffer Z (1.28X Transcription Buffer, 12.82 mM MgCl_2_, 0.05% Tween 20) for ZTP riboswitch assays or 18.75 μl per sample volume of Elution Buffer G (1.33X Transcription Buffer, 13.33 mM MgCl_2_, 0.07% Tween 20, 0.67 μM NusA) for ppGpp riboswitch assays. For ZTP riboswitch assays, eluted ^C3-SC1^TECs were pre-warmed at 37 *°*C for 2 minutes before 19.5 μl of ^C3-SC1^TECs were mixed with 5.5 μl of pre-warmed Chase Mix Z (2.5 μl 5 mM NTPs, 2.5 μl 100 μg/ml rifampicin, 0.5 μl of ZMP (Sigma Aldrich) in DMSO at 50X final ZMP concentration), incubated at 37 *°*C for 5 minutes, and mixed with 125 μl of Stop Solution. For ppGpp riboswitch assays, eluted ^C3-SC1^TECs were pre-warmed at 37 *°*C for 2 minutes before 18.75 μl of ^C3-SC1^TECs were mixed with 6.25 μl of pre-warmed Chase Mix G (2.5 μl 5 mM NTPs, 2.5 μl 100 μg/ml rifampicin, 1.25 μl of ppGpp (Jena Bioscience) in nuclease-free water at 20X final ppGpp concentration), incubated at 37 *°*C for 5 minutes, and mixed with 125 μl of Stop Solution.

Transcription termination assays for leader-less riboswitches were performed as follows: 25 μl transcription reactions containing 1X Transcription Buffer, 0.1 mg/ml BSA, 10 mM MgCl_2_, 1X AGU Start NTPs (2.5 μM ATP, 2.5 μM GTP, 1.5 μM UTP), 0.2 μCi/μl [ -^32^P]UTP, 5 nM DNA template 3 or 5 (Tables S2 and S3), and 0.016 U/μl *E. coli* RNAP were prepared in a 1.7 ml microcentrifuge tube on ice; at this point the total reaction volume was 19.5 μl per sample volume for ZTP riboswitch assays and 18.75 μl per sample volume for ppGpp riboswitch assays due to the omission of Chase Mix Z and Chase Mix G, respectively. Transcription reactions were incubated at 37 *°*C for 10 minutes to walk RNAP to the first C in the riboswitch sequence (+15 for the ZTP riboswitch and +17 for the ppGpp riboswitch). For ZTP riboswitch assays, 19.5 μl of the master mix was mixed with 5.5 μl of pre-warmed Chase Mix Z, incubated at 37 *°*C for 5 minutes, and mixed with 125 μl of Stop Solution. For ppGpp riboswitch assays, 18.75 μl of the master mix was mixed with 6.25 μl of pre-warmed Chase Mix G, incubated at 37 *°*C for 5 minutes, and mixed with 125 μl of Stop Solution. The samples were processed as described below in the section *Purification, sequencing gel electrophoresis, and detection of radiolabeled RNA*.

### Purification, sequencing gel electrophoresis, and detection of radiolabeled RNA

Radiolabeled RNA from *in vitro* transcription reactions was processed and analyzed exactly as described previously (35). Briefly, 150 μl samples (25 μl sample + 125 μl Stop Solution) were mixed with an equal volume (150 μl) of Tris (pH 8) buffered phenol:chloroform:isoamyl alcohol (25:24:1, v/v), mixed by vortexing and inversion, and centrifuged at 18,500 × g and 4 *°*C for 5 minutes. The aqueous phase was collected and transferred to a new tube. Nucleic acids were precipitated by adding 3 sample volumes (450 μl) of 100% ethanol and 1 or 1.2 μl of GlycoBlue Coprecipitant and chilling at -70 *°*C for at least 30 minutes. The samples were centrifuged at 18,500 × g and 4 *°*C for 30 minutes, the supernatant was removed, the samples were centrifuged again briefly to pull down residual ethanol, and residual ethanol was removed. The pellets were dissolved in 6.5 μl of Formamide Loading Dye, denatured by heating at 95 *°*C for 5 minutes, loaded on a pre-warmed 7.5 M urea, 12% polyacrylamide, 35 × 43 cm, 0.4 mm thick sequencing gel in a Model S2 Sequencer Apparatus, and run at 1400 V for 2.5 to 3 h. The resulting gel was exposed to a storage phosphor screen for 12 to 16 h, and the storage phosphor screen was scanned using an Amersham Typhoon IP Biomolecular Imager (Cytiva).

### Quantification

Quantification of band intensity was performed by analyzing gel image TIFF files using ImageJ 1.53k exactly as described previously (34). The distribution of DNA between pellet and supernatant fractions (Figure 2), the purity of ^C3-SC1^TECs (Figure 3B), and the distribution of input DNA between S1, W, P, and S2 fractions (Figure 3C) were calculated exactly as described previously for roadblocked TECs (34).

In experiments that used radiolabeled ^C3-SC1^TECs, all [ -^32^P]UTP was removed before the ^C3-SC1^TECs were chased; therefore all RNA species were uniformly labeled and no normalization was applied. In functional assays using leader-less riboswitches, transcripts were considered end-labeled, as described previously (8), due to the high probability of [ -^32^P]UTP incorporation during the initial walk (∼4.25% per U nucleotide) and the low probability of [ -^32^P]UTP incorporation during the chase (∼0.013% per U nucleotide) and no normalization was applied. In Figure 4B, fraction full length was calculated by dividing the band intensity of etheno-dA stalled transcripts by the total band intensity of stalled and synched transcripts. In Figures 5B and 5C, the fraction of synched/stalled TECs that were retained on beads was calculated by dividing the band intensity of RNA in the pellet fraction by the sum of pellet and supernatant band intensities for each indicated complex. In Figure 5D, the fraction of TECs chased was calculated by dividing the sum of the pellet and supernatant band intensities of etheno-dA stalled transcripts by the total band intensity of stalled and synched transcripts in both fractions. In Figures 6B and 6C, the fraction of terminator readthrough was calculated by dividing the band intensity of readthrough transcripts by the total band intensity of terminated and readthrough transcripts.

### Reproducibility of the method

>10 independent preparations of ^C3-SC1^TECs were performed by S.L.K. and C.E.S. using separate reagent stocks and assessed by EMSA by using the final protocol (or a closely related protocol that yielded indistinguishable complexes). Examples of these data are shown in Figure 3 and Supplementary Figure S1A.

## Data availability

All data are contained in the manuscript as plotted values or representative gels. Source files in TIFF and/or gel format are available from the corresponding author (E.J.S.) upon request.

## Supporting information

This article contains supporting information.

## Author Contributions

E.J.S. conceptualization; S.L.K., C.E.S., and E.J.S. formal analysis; E.J.S. funding acquisition; S.L.K., C.E.S., and E.J.S. investigation; S.L.K., C.E.S., and E.J.S. methodology; E.J.S. supervision; S.L.K., C.E.S. validation; S.L.K., C.E.S., and E.J.S. writing-original draft; S.L.K., C.E.S., and E.J.S. writing-review and editing.

## Funding

This work was supported by startup funding from the University at Buffalo (to E.J.S.).

## Conflict of Interest

The authors have no conflicts of interest with the contents of this article.

## Abbreviations

The abbreviations used are

RNAP: RNA polymerase
TEC: transcription elongation complex
C3: capture sequence 3
SC1: structure cassette 1
etheno-dA: 1,N^6^-etheno-2’deoxyadenosine
ZMP: 5-aminoimidazole-4-carboxamide-1-β-D-ribofuranosyl 5’-monophosphate
ppGpp: guanosine-3’,5’-bisdiphosphate
TBE: tris-borate-EDTA
P: pellet
S: supernatant
T: terminated
RT: readthrough

## Materials Included

**Figure S1.**
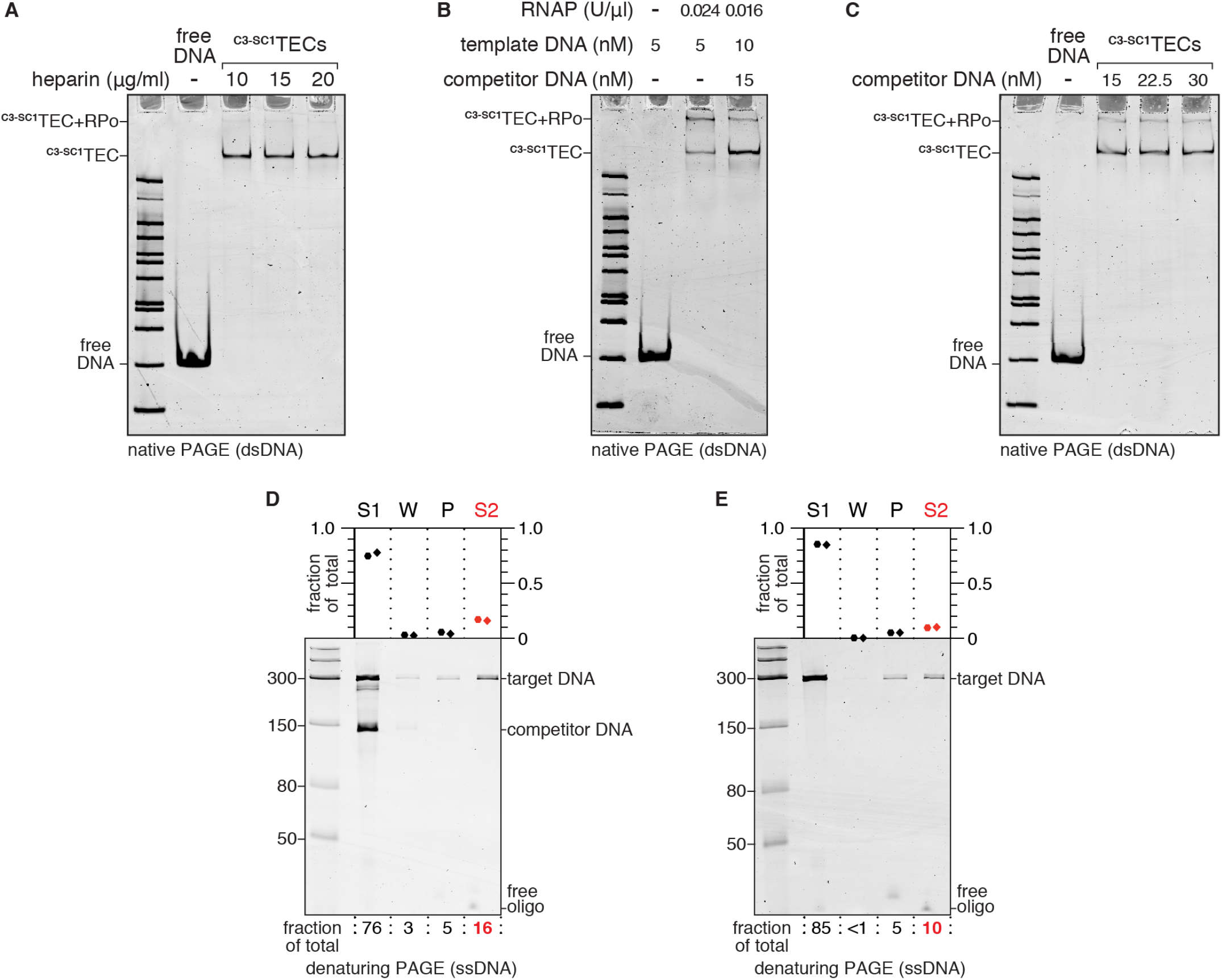
Alternative conditions for ^C3-SC1^TEC purification. *A*, EMSA of ^C3-SC1^TECs that were purified using variable amounts of heparin. *B*, EMSA of ^C3-SC1^TECs that were purified using a competitor DNA template instead of heparin. Omitting the competitor DNA and including excess RNAP favored the formation of slow-migrating complexes that contain both a ^C3-SC1^TEC and an open promoter complex. *C*, EMSA of ^C3-SC1^TECs that were purified using variable amounts of competitor DNA. *D*, denaturing PAGE analysis of purification fractions for ^C3-SC1^TECs that were prepared using 15 nM competitor DNA. *E*, denaturing PAGE analysis of purification fractions for ^C3-SC1^TECs that were prepared using 20 μg/ml heparin but without rifampicin.

**Figure S2.**
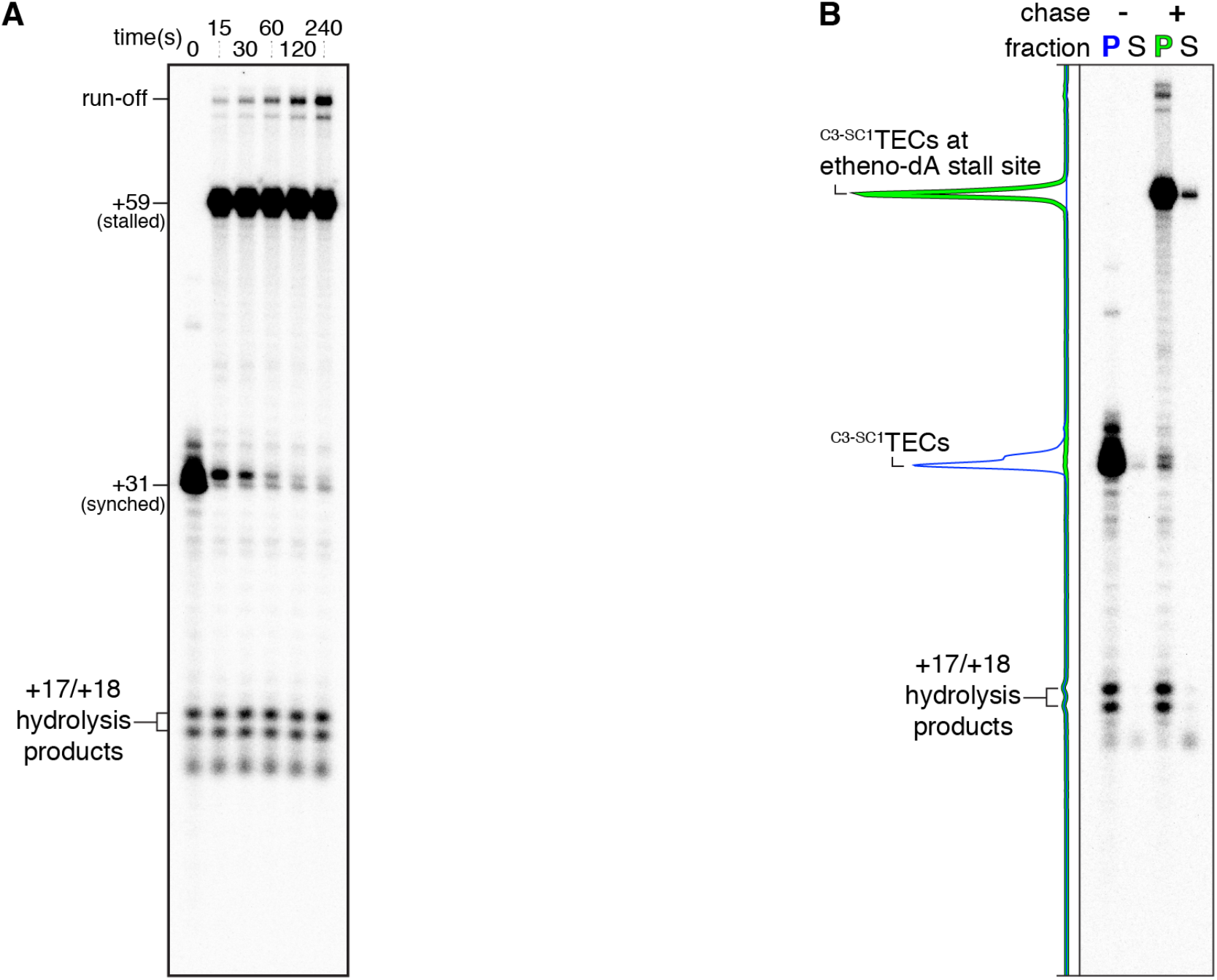
Additional visualization of gels for ^C3-SC1^TEC kinetics and solid-phase transcription assays. Gels from Figure 4A (*A*) and Figure 5A (*B*) are presented here with a darker grayscale setting to visualize trace RNAs.

**Table S1.**
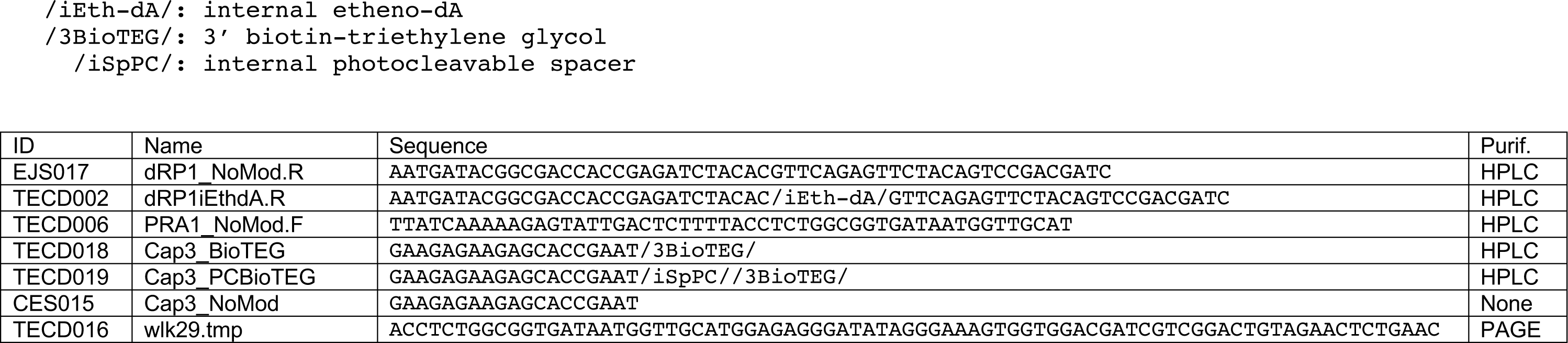
Oligonucleotides used in this study. Below is a table of oligonucleotides used in this study. The modification codes defined below are used for compatibility with Integrated DNA Technologies ordering. DNA containing internal etheno-dA requires an off-catalog order.

**Table S2.**
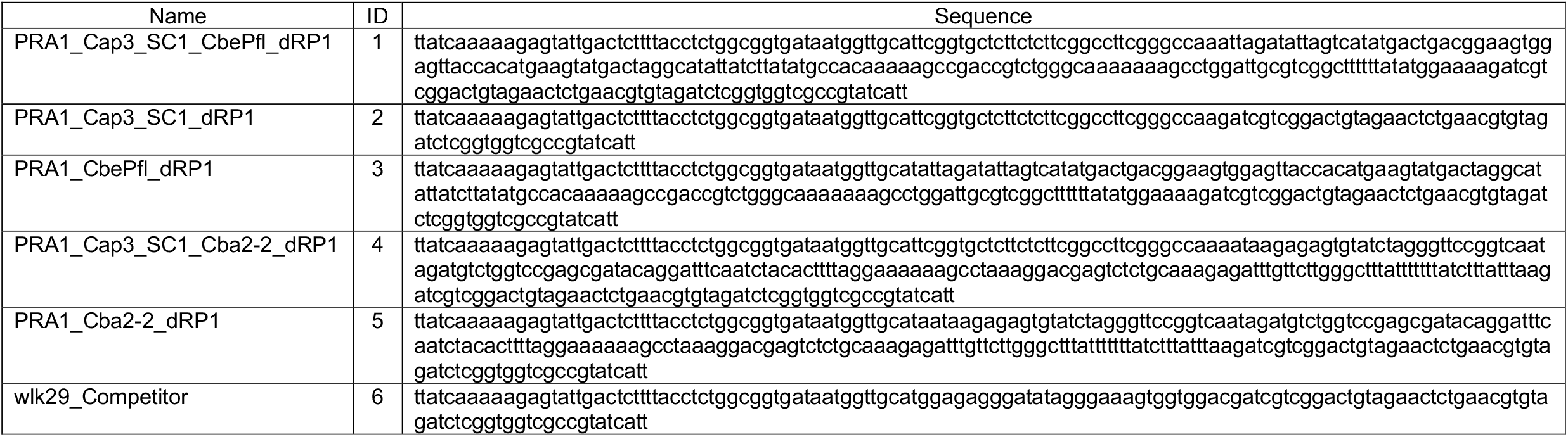
DNA template sequences. Below is a table of containing the sequence of each DNA template. Fully annotated versions are available at Benchling (See ‘DNA Template Sequences’ in Experimental Procedures for hyperlinks).

**Table S3.** DNA templates prepared for this study. Below is a table of DNA templates that were prepared for this study, including the primers, plasmids, and template oligos used, DNA modifications, the PCR polymerase used, whether translesion synthesis was performed, which reaction clean up protocol was used (see Experimental Procedures), and the figures in which each DNA template was used.

| ID | Fwd Primer | Rev Primer | Template | Modifications | PCR Polymerase | Translesion Synthesis | Clean Up | Used in Fig(s) |
| --- | --- | --- | --- | --- | --- | --- | --- | --- |
| 1 | TECD006 | EJS017 | pCES002 | none | Vent (exo-) | N/A | Gel extracted | 2, 3B, 3C, 6B, 6C, S1A, S1B, S1C, S1D, S1E |
| 2 | TECD006 | TECD002 | Gel-purified linear DNA from pCES003 | Int etheno-dA | Q5 | Yes | PCR clean-up | 4, 5, S2A, S2B |
| 3 | TECD006 | EJS017 | pCES004 | none | Vent (exo-) | N/A | Gel extraction | 6B, 6C |
| 4 | TECD006 | EJS017 | pCES005 | none | Vent (exo-) | N/A | Gel extraction | 6E, 6F |
| 5 | TECD006 | EJS017 | pCES006 | none | Vent (exo-) | N/A | Gel extraction | 6E, 6F |
| 6 | TECD006 | TECD002 | TECD016 | Int etheno-dA | Q5 | Yes | PCR clean-up | S1B, S1C, S1D |

## Notes

### Competing Interest Statement

The authors have declared no competing interest.

## References

1. Watters, K. E., Strobel, E. J., Yu, A. M., Lis, J. T., and Lucks, J. B. (2016) Cotranscriptional folding of a riboswitch at nucleotide resolution. Nat Struct Mol Biol 23, 1124–1131

2. Strobel, E. J., Watters, K. E., Nedialkov, Y., Artsimovitch, I., and Lucks, J. B. (2017) Distributed biotin-streptavidin transcription roadblocks for mapping cotranscriptional RNA folding. Nucleic Acids Res 45, e109

3. Frieda, K. L., and Block, S. M. (2012) Direct observation of cotranscriptional folding in an adenine riboswitch. Science 338, 397–400

4. Buenrostro, J. D., Araya, C. L., Chircus, L. M., Layton, C. J., Chang, H. Y., Snyder, M. P., and Greenleaf, W. J. (2014) Quantitative analysis of RNA-protein interactions on a massively parallel array reveals biophysical and evolutionary landscapes. Nat Biotechnol 32, 562–568

5. Denny, S. K., Bisaria, N., Yesselman, J. D., Das, R., Herschlag, D., and Greenleaf, W. J. (2018) High-Throughput Investigation of Diverse Junction Elements in RNA Tertiary Folding. Cell 174, 377–390 e320

6. Widom, J. R., Nedialkov, Y. A., Rai, V., Hayes, R. L., Brooks, C. L., 3rd, Artsimovitch, I., and Walter, N. G. (2018) Ligand Modulates Cross-Coupling between Riboswitch Folding and Transcriptional Pausing. Mol Cell 72, 541–552 e546

7. Denny, S. K., and Greenleaf, W. J. (2019) Linking RNA Sequence, Structure, and Function on Massively Parallel High-Throughput Sequencers. Cold Spring Harb Perspect Biol 11

8. Strobel, E. J., Cheng, L., Berman, K. E., Carlson, P. D., and Lucks, J. B. (2019) A ligand-gated strand displacement mechanism for ZTP riboswitch transcription control. Nat Chem Biol 15, 1067–1076

9. Fukuda, S., Yan, S., Komi, Y., Sun, M., Gabizon, R., and Bustamante, C. (2020) The Biogenesis of SRP RNA Is Modulated by an RNA Folding Intermediate Attained during Transcription. Mol Cell 77, 241–250 e248

10. Chatterjee, S., Chauvier, A., Dandpat, S. S., Artsimovitch, I., and Walter, N. G. (2021) A translational riboswitch coordinates nascent transcription-translation coupling. Proc Natl Acad Sci U S A 118

11. Chauvier, A., Ajmera, P., Yadav, R., and Walter, N. G. (2021) Dynamic competition between a ligand and transcription factor NusA governs riboswitch-mediated transcription regulation. Proc Natl Acad Sci U S A 118

12. Chauvier, A., St-Pierre, P., Nadon, J. F., Hien, E. D. M., Perez-Gonzalez, C., Eschbach, S. H., Lamontagne, A. M., Penedo, J. C., and Lafontaine, D. A. (2021) Monitoring RNA dynamics in native transcriptional complexes. Proc Natl Acad Sci U S A 118

13. Landick, R., Wang, D., and Chan, C. L. (1996) Quantitative analysis of transcriptional pausing by Escherichia coli RNA polymerase: his leader pause site as paradigm. Methods Enzymol 274, 334–353

14. Levin, J. R., Krummel, B., and Chamberlin, M. J. (1987) Isolation and properties of transcribing ternary complexes of Escherichia coli RNA polymerase positioned at a single template base. J Mol Biol 196, 85–100

15. Chan, C. L., and Landick, R. (1989) The Salmonella typhimurium his operon leader region contains an RNA hairpin-dependent transcription pause site. Mechanistic implications of the effect on pausing of altered RNA hairpins. J Biol Chem 264, 20796–20804

16. Olejnik, J., Sonar, S., Krzymanska-Olejnik, E., and Rothschild, K. J. (1995) Photocleavable biotin derivatives: a versatile approach for the isolation of biomolecules. Proc Natl Acad Sci U S A 92, 7590–7594

17. Merino, E. J., Wilkinson, K. A., Coughlan, J. L., and Weeks, K. M. (2005) RNA structure analysis at single nucleotide resolution by selective 2’-hydroxyl acylation and primer extension (SHAPE). J Am Chem Soc 127, 4223–4231

18. Wilkinson, K. A., Merino, E. J., and Weeks, K. M. (2006) Selective 2’-hydroxyl acylation analyzed by primer extension (SHAPE): quantitative RNA structure analysis at single nucleotide resolution. Nat Protoc 1, 1610–1616

19. Hein, P. P., Palangat, M., and Landick, R. (2011) RNA transcript 3’-proximal sequence affects translocation bias of RNA polymerase. Biochemistry 50, 7002–7014

20. Larson, M. H., Mooney, R. A., Peters, J. M., Windgassen, T., Nayak, D., Gross, C. A., Block, S. M., Greenleaf, W. J., Landick, R., and Weissman, J. S. (2014) A pause sequence enriched at translation start sites drives transcription dynamics in vivo. Science 344, 1042–1047

21. Vvedenskaya, I. O., Vahedian-Movahed, H., Bird, J. G., Knoblauch, J. G., Goldman, S. R., Zhang, Y., Ebright, R. H., and Nickels, B. E. (2014) Interactions between RNA polymerase and the “core recognition element” counteract pausing. Science 344, 1285–1289

22. Saba, J., Chua, X. Y., Mishanina, T. V., Nayak, D., Windgassen, T. A., Mooney, R. A., and Landick, R. (2019) The elemental mechanism of transcriptional pausing. Elife 8

23. Winkelman, J. T., Pukhrambam, C., Vvedenskaya, I. O., Zhang, Y., Taylor, D. M., Shah, P., Ebright, R. H., and Nickels, B. E. (2020) XACT-Seq Comprehensively Defines the Promoter-Position and Promoter-Sequence Determinants for Initial-Transcription Pausing. Mol Cell 79, 797–811 e798

24. Heyduk, E., and Heyduk, T. (2018) DNA template sequence control of bacterial RNA polymerase escape from the promoter. Nucleic Acids Res 46, 4469–4486

25. Jacques, J. P., and Kolakofsky, D. (1991) Pseudo-templated transcription in prokaryotic and eukaryotic organisms. Genes Dev 5, 707–713

26. Turnbough, C. L., Jr., and Switzer, R. L. (2008) Regulation of pyrimidine biosynthetic gene expression in bacteria: repression without repressors. Microbiol Mol Biol Rev 72, 266–300, table of contents

27. Murakami, K. S., Shin, Y., Turnbough, C. L., Jr., and Molodtsov, V. (2017) X-ray crystal structure of a reiterative transcription complex reveals an atypical RNA extension pathway. Proc Natl Acad Sci U S A 114, 8211–8216

28. Liu, Y., Winkelman, J. T., Yu, L., Pukhrambam, C., Zhang, Y., Nickels, B. E., and Ebright, R. H. (2021) Structural and mechanistic basis of RNA extension in reiterative transcription initiation: RNA slipping without DNA scrunching. bioRxiv

29. Reuter, J. S., and Mathews, D. H. (2010) RNAstructure: software for RNA secondary structure prediction and analysis. BMC Bioinformatics 11, 129

30. Komissarova, N., and Kashlev, M. (1998) Functional topography of nascent RNA in elongation intermediates of RNA polymerase. Proc Natl Acad Sci U S A 95, 14699–14704

31. Komissarova, N., and Kashlev, M. (1997) Transcriptional arrest: Escherichia coli RNA polymerase translocates backward, leaving the 3’ end of the RNA intact and extruded. Proc Natl Acad Sci U S A 94, 1755–1760

32. Artsimovitch, I., and Landick, R. (2000) Pausing by bacterial RNA polymerase is mediated by mechanistically distinct classes of signals. Proc Natl Acad Sci U S A 97, 7090–7095

33. Zhang, J., and Landick, R. (2016) A Two-Way Street: Regulatory Interplay between RNA Polymerase and Nascent RNA Structure. Trends Biochem Sci 41, 293–310

34. Strobel, E. J. (2021) Preparation of E. coli RNA polymerase transcription elongation complexes by selective photoelution from magnetic beads. J Biol Chem 297, 100812

35. Strobel, E. J., Lis, J. T., and Lucks, J. B. (2020) Chemical roadblocking of DNA transcription for nascent RNA display. J Biol Chem 295, 6401–6412

36. Pupov, D., Ignatov, A., Agapov, A., and Kulbachinskiy, A. (2019) Distinct effects of DNA lesions on RNA synthesis by Escherichia coli RNA polymerase. Biochem Biophys Res Commun 510, 122–127

37. Kim, P. B., Nelson, J. W., and Breaker, R. R. (2015) An ancient riboswitch class in bacteria regulates purine biosynthesis and one-carbon metabolism. Mol Cell 57, 317–328

38. Sherlock, M. E., Sudarsan, N., and Breaker, R. R. (2018) Riboswitches for the alarmone ppGpp expand the collection of RNA-based signaling systems. Proc Natl Acad Sci U S A 115, 6052–6057

39. Henderson, K. L., Felth, L. C., Molzahn, C. M., Shkel, I., Wang, S., Chhabra, M., Ruff, E. F., Bieter, L., Kraft, J. E., and Record, M. T., Jr. (2017) Mechanism of transcription initiation and promoter escape by E. coli RNA polymerase. Proc Natl Acad Sci U S A 114, E3032–E3040

40. Sidorenkov, I., Komissarova, N., and Kashlev, M. (1998) Crucial role of the RNA:DNA hybrid in the processivity of transcription. Mol Cell 2, 55–64

41. Komissarova, N., Kireeva, M. L., Becker, J., Sidorenkov, I., and Kashlev, M. (2003) Engineering of elongation complexes of bacterial and yeast RNA polymerases. Methods Enzymol 371, 233–251

42. Drogalis, L. K., and Batey, R. T. (2020) Requirements for efficient ligand-gated co-transcriptional switching in designed variants of the B. subtilis pbuE adenine-responsive riboswitch in E. coli. PLoS One 15, e0243155

43. Strobel, E. J. (2021) Preparation and Characterization of Internally Modified DNA Templates for Chemical Transcription Roadblocking. Bio Protoc 11, e4141

